# Neural predictors of subjective fear depend on the situation

**DOI:** 10.1101/2022.10.20.513114

**Authors:** Yiyu Wang, Philip A. Kragel, Ajay B. Satpute

**Affiliations:** Department of Psychology, Northeastern University; Department of Psychology, Emory University; Department of Radiology, Massachusetts General Hospital

**Keywords:** Fear, subjective experience, multivariate pattern analysis

## Abstract

The extent to which neural representations of fear experience depend on or generalize across the situational context has remained unclear. We systematically manipulated variation within and across three distinct fear-evocative situations including fear of heights, spiders, and social threats. Participants (n=21, 10 females and 11 males) viewed 20 second clips depicting spiders, heights, or social encounters, and rated fear after each video. Searchlight multivoxel pattern analysis (MVPA) was used to identify whether and which brain regions carry information that predicts fear experience, and the degree to which the fear-predictive neural codes in these areas depend upon or generalize across the situations. The overwhelming majority of brain regions carrying information about fear did so in a situation dependent manner. These findings suggest that local neural representations of fear experience are unlikely to involve a singular pattern, but rather a collection of multiple heterogeneous brain states

---

For over a century, philosophers, psychologists, and neuroscientists have debated the nature of emotions (Barrett & Satpute, 2019; Dalgleish et al., 2009; Gendron & Feldman Barrett, 2009). Much of this debate concerns the uniformity or heterogeneity of representation for particular emotion categories (Lindquist et al., 2013; Mobbs et al., 2019). For example, is there a particular brain state that underlies fearful experiences (Celeghin et al., 2017; Nummenmaa & Saarimäki, 2019; Vytal & Hamann, 2010), or does fear involve a collection of heterogeneous brain states (Doyle et al., 2022; Satpute Lindquist, 2019; Wilson-Mendenhall et al., 2011)? Addressing this question has broad implications for translational neuroscience models of mood and anxiety disorders (Fanselow & Pennington, 2017; LeDoux Pine, 2016). Of relevance to this question are recent functional magnetic resonance imaging (fMRI) studies that have searched for the brain “signatures” of emotion categories (Kassam et al., 2013; Kragel & LaBar, 2015, 2016; Saarimäki et al., 2016, 2018). Brain signatures (sometimes referred to as “neuromarkers” or “neural signatures”) are a type of multivoxel pattern analysis (MVPA) that use brain data to predict the affective state of an individual (Kragel et al., 2018). Several studies have shown that brain signatures of emotion that are optimal for prediction draw on information that is widely distributed throughout much of the brain. However, the degree to which this information is organized into a single prototypical pattern or reflects an amalgam of heterogeneous functional states remains unclear because neural signatures do not inherently provide direct insight about neural representation (Kragel et al., 2018; Lindquist et al., 2022). Significant classification is possible even if few, or even none, of the brain regions individually carry category-level representations of emotion (Azari et al., 2020; Clark-Polner et al., 2017; Kragel et al., 2018; Lindquist et al., 2022). Alternatively, a handful of univariate fMRI studies have shown that distinct sets of brain regions are engaged depending on the fear-evocative content (e.g. pictures of spiders, blood/injections/injuries, social encounters) (Caseras et al., 2010; Lueken et al., 2011; Michalowski et al., 2017). These findings are suggestive of heterogeneity in the neural representation of fear based on situated content. However, two limitations to this work preclude a clear conclusion. First, to investigate fear, these studies either compared categories of stimuli with distinct semantic content (e.g. stereotypically fearful stimuli, e.g., spiders or snakes, vs. “neutral” stimuli, e.g., ordinary objects), participant groups (phobic v. non-phobic; (Caseras et al., 2010), or both (Lueken et al., 2011; Michalowski et al., 2017). Accordingly, neural differences may be due to the semantic content of the stimuli or between individuals that are unrelated to fear. Second, these studies focused on activation magnitude, yet patterns of activation may carry information about psychological states irrespective of differences in activation magnitude. Here, we tested the degree to which functional activity in any given brain region predicts fear in a situation-general or situation-dependent manner. Participants viewed 20 second clips depicting spiders, heights, or social encounters, and rated fear after each video. We selected these situations because they span a wide variety of properties. For instance, while fear is often studied in a predator-prey context, fear of heights is potent and yet does not involve a predator. Critically, video stimuli were also curated to evoke a wide range of fear within each situation (Figure 1). This design enabled us to systematically examine the neural predictors of fear within and across each situation. We then used searchlight MVPA (Kriegeskorte et al., 2006) with LASSO-PCR (Tibshirani, 1996; Wager et al., 2011) to identify brain regions with functional information that predicts fear ratings. A brain region may carry fear-predictive information that generalizes across all situations using the same neural codes (i.e., situation-general, shared model parameters) or using different neural codes (i.e., situation-general, unshared model parameters), or situation-dependent information. Thus, we trained our models in two distinct ways to test these possibilities.

**Figure 1.**
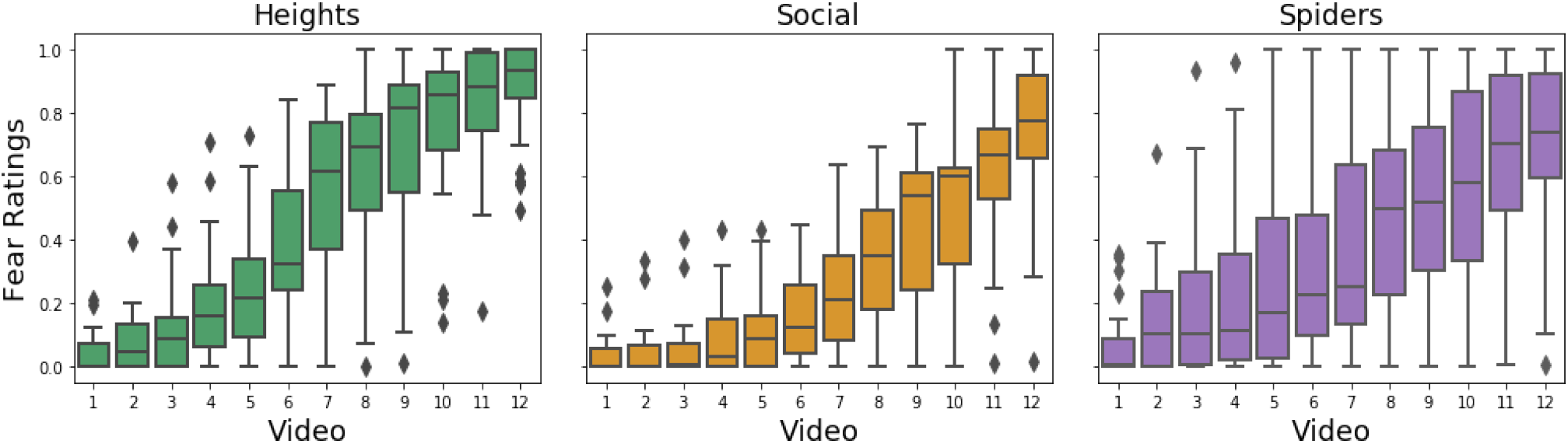
Within Category Variation in Fear Ratings. Each box and whiskers plot shows the mean and variation (quartiles and max/min values) in fear ratings (0 “low fear”, 1 “high fear”) across participants by video rank. Videos (x-axis) are sorted from lowest to highest on subjective fear ratings by participant, regardless of the video identity, since different videos evoked different levels of fear experience across participants. The plot illustrates that the videos were effective at evoking a wide range of fear experiences within each situation and for each person. Thus, MVPA can be used to examine which brain regions predict fear within and across situations.

## Methods

### Participants

Neurotypical participants who reported no clinical psychiatric diagnosis were recruited from the greater Los Angeles area. Exclusion criteria consisted of claustrophobia, psychiatric medication, left-handedness, metal in the body, and age (under 18 years or over 55 years). After excluding three individuals with excessive motion (criteria described below), the effective sample included 21 participants (11 male, 10 female, ages 22-40, mean age = 30.4).

### Stimuli

Thirty six, silent videos were used in the experiment (12 videos per situation; duration: 18-22 s/video). While silent videos might be less evocative, they provide a more conservative test of the situation-dependence hypothesis, since it has already been shown that neural responses during affective experiences are modality dependent. For ecological validity, all videos depicted naturalistic footage and were shot from an immersive 1st person perspective. Videos were selected to be relatively stable (i.e., did not involve dramatic changes or “jump scares”) to mitigate motion artifacts and maintain the consistency of psychological experience across the duration. Video stimuli were obtained and normed in an independent online sample (see Extended Data: Stimulus Norms). Stimuli were curated to elicit a wide range of variation in self-reported fear across three distinct situations such that models could be estimated to predict fear within each situation (Figure 1). For example, a heights video normatively high in fear ratings depicts first-person footage of walking along the edge of a sheer cliff, whereas one normatively low in fear depicts first-person footage of walking down a set of stairs. While norms were used to select stimuli for inclusion, analyses were conducted using subjective reports. A short description of the content and the normative ratings from the independent online sample for each video are available online: https://github.com/yiyuwang/AffVids_mvpa/tree/main/video_info.

### Experimental Task

Video stimuli were presented across three functional runs (12 videos/run) in the MRI scanner. Each run had an equal number of videos from each situation category, and each run had an equal number of normatively high and low fear (based on median splits of normative ratings) videos within each situation (see Extended Data Figure 1-1). The order of video stimulus presentations was pseudo randomized to ensure uniformity of stimulus types over time. First, videos of a given category were preceded by videos of the same and different categories equally often. Second, videos with a given normative fear rating were preceded by videos of higher and lower normative fear ratings equally often. Participants were instructed, and reminded in between scans, “to immerse yourself in the situation shown”, and also to “respond according to how you, in particular, feel in response to viewing the videos”. Immediately after each 20 s video, participants made self-report ratings on a sliding scale for experienced fear, arousal, and valence. Fear and arousal were measured on a scale ranging from “low” to “high”, and valence was measured on a scale ranging from “negative” to “positive”. Participants used a trackball to move a cursor along a continuous scale and then clicked a button under their right thumb to log their rating. Four seconds were allotted to make each rating (12 seconds total). The task also included an anticipatory period before each video, wherein the word (“Heights”, “Social”, or “Spider”) corresponding to the category of the upcoming video was presented for 3 seconds, followed by a jittered fixation interval of 3-5 seconds, during which participants rated their expected fear on a sliding scale anchored by “low” to “high”. The purpose of this period was to mitigate effects pertaining to semantic updating that would otherwise occur when transitioning from a fixation cross to a rich visual image, and to address other research questions regarding anticipatory activity prior to video watching. This period was not analyzed to address the present hypotheses. Trials were presented across three, 9 minute runs. Participants failed to provide fear ratings within the allotted time on a small proportion of trials. Missing fear ratings were interpolated (see Extended Data: Interpolation) and included in analyses. Stimuli were presented using MATLAB (MathWorks) and the Psychophysics Toolbox, and behavioral responses were recorded using a scanner-compatible trackball.

### fMRI data acquisition and preprocessing

MRI data were collected using a 3T Siemens Trio MRI scanner. Functional images were acquired in interleaved order using a T2*-weighted multiband echo planar imaging (EPI) pulse sequence (transverse slices, TR=1000ms, TE = 3000ms, flip angle=60°, FOV = 200mm, 2.5 mm thickness slices, voxel dimension= 2.5 x 2.5 x 2.5 mm, Phase Encoding Direction anterior to posterior (AP), multi band acceleration factor = 4). Functional scans included coverage of amygdala and orbitofrontal cortex (see Extended Data Figures 3-3 and 3-4). Anatomical images were acquired at the start of the session with a T1-weighted pulse sequence (TR = 2400ms, TE = 2600ms, flip angle=8°, FOV = 256mm, 1 mm thickness slices, voxel dimension= 1 x 1 x 1 mm). Image volumes were preprocessed using fMRIprep (Esteban et al., 2019). Preprocessing included motion correction, slice-timing correction, removal of high frequency drifts using a temporal high-pass filter (discrete cosine transform, 100s cutoff), and spatial smoothing (6mm FWHM). For all analyses, functional volumes were downsampled to a 3 mm space to speed searchlight analyses and registered to participants’ anatomical image and then to a standard template (MNI152) using FSL FLIRT (Jenkinson et al., 2002). Participants with at least two runs without excessive head motion (defined as > 2 mm maximum framewise displacement), were included in the analysis yielding 18 participants with three runs of data and 3 participants with two runs of data.

### General Linear Model

A general linear model (GLM) was used to model the neural data. The GLM included a separate boxcar regressor for each video stimulus, convolved with a canonical hemodynamic response function from SPM12. Nuisance regressors included six regressors corresponding with motion parameters, three regressors for physiological noise artifacts (CSF, white matter, framewise displacement), and non-steady states outliers (stick function per outlier). Three regressors were included to model low-level visual properties of the stimuli. Specifically, luminance, contrast, and the complexity of each extracted frame were calculated using Matlab scripts at https://github.com/yiyuwang/AffVids_mvpa/tree/main/calculate_visual_property. We extracted pixel values from the frame occuring at the beginning of each TR. Luminance was calculated as the mean value of the grayscale image of the frame. Contrast was calculated as the difference between the maximum luminance and the minimum luminance. Complexity was calculated as the entropy of the grayscale image of the frame. The GLM was conducted using custom scripts in the Python nilearn module. Beta maps for each video and participant were used for training the searchlight LASSO-PCR (see below).

### Searchlight LASSO-PCR

We analyzed the data using a searchlight multivariate pattern analysis approach. Betas from the GLM were extracted from voxels within the voxel’s searchlight neighborhood using a 15mm (5 voxel) radius. Because voxel data is non-independent, we first run a principal component analysis (PCA) with the same number of components as the number voxels in the searchlight. The PCA transforms non-independent activity across voxels as a set of orthogonal components. The components are then used as regressors in a Least Absolute Shrinkage and Selection Operator (LASSO) Regression (Chang et al., 2015; Tibshirani, 1996) to predict continuous fear ratings. A relatively lenient penalty term of 0.05 was used since the searchlight analysis already constrains the dimensionality of the fMRI data. The analysis was performed using modified functions from the scikit-learn and nilearn Python module (Pedregosa et al., 2011). All codes are publicly available at https://github.com/yiyuwang/AffVids_mvpa.

### Cross Validation

Prior studies that examined the neural predictors of fear trained and tested their model across groups of individuals (Kragel & LaBar, 2015; Zhou et al., 2021). Here, we follow suit with this approach by combining data across participants and training and testing our models using three-fold, leave-whole-subject-out, cross validation to increase robustness (Poldrack et al., 2020). Participants were randomly divided into three groups (folds) of seven participants each (we selected three since it evenly divides the 21 participants in the sample). Models (i.e. searchlight with LASSO-PCR) were iteratively trained on two groups and tested on the left-out group. The dot product of the model weights from LASSO-PCR from the training data and activation data from the testing sample yields predicted fear ratings. Pearson correlations between the predicted fear ratings and the actual fear ratings were calculated and assigned to the center voxel of the spherical searchlight. After iterating the searchlight across the whole brain, the analysis resulted in a whole brain map of the correlation values between the predicted ratings and the observed ratings of the testing sample for each of the three folds. We averaged the maps from the three folds as our final result.

### Model Training and Testing

In the across-situation training method, models were trained on data across all three stimulus categories. In each fold, the model was trained on data corresponding with all thirty-six videos from fourteen participants (i.e., 504 samples), and was then tested on data for each stimulus category from the seven left out participants. In the situation-by-situation training method, models were trained on data from one stimulus category at a time. For example, the model was trained using data corresponding with twelve heights video stimuli from fourteen participants (i.e., 168 samples), and then tested on data for heights video stimuli from the seven left-out participants. The across-situation training method has the advantage of more training samples, which yields more robust results. Thus, we performed analyses with more balanced training sets, too, and found that this training advantage for the situation-general model is unlikely to impact our conclusions (Extended data, Figure 3-2).

### Permutation Testing and Statistical Correction

Permutation testing (N = 1,200/voxel-wise neighborhood) was used to identify voxels with nonzero predictions. Models were trained and tested using shuffled data to generate a null distribution of correlation values. Comparing the original correlation values against the null distribution resulted in a statistical map that could be thresholded, using the p-values obtained for each voxel. A Familywise Error (FWE) rate of 0.05 was used to threshold the permutation test (Nichols & Holmes, 2002; Nichols & Hayasaka, 2003).

## Results

Behavioral findings confirmed the central aim of the task design, namely, that fear ratings varied from low to high levels within each content condition and across participants (Figure 1). Thus, we proceeded to examine whether and which brain regions contained functional activity that predicts fear ratings within and across situations. There are three possible outcomes. First, population activity of neurons in a brain region may code for fear in the same way across situations. If so, then functional activity for a given brain region may predict fear ratings across situations using the same “neural codes” (i.e., model parameters). Second, population activity may code for fear in different ways for different situations, for example, if a brain region contains segregated neural pathways or the same pathway organizes into distinct configurations. If so, then functional activity for a given brain region may predict fear ratings across situations but requires different neural codes. These hypothetical possibilities dictate two different model training approaches. If a brain region carries situation general neural codes, model training should include instances across situations to enable the model to best learn which signals are indeed situation general. Alternatively, if a brain region carries situation dependent neural activity, model training should occur situation by situation. Thus, we trained our models using both approaches to assess whether the conclusions depend on the theoretical perspective and affiliated analytical approach. Finally, a third possibility is that population activity may code for fear in one or two situations, but not all three, suggesting that both the brain regions and neural codes that predict fear are situation dependent. Thus, using both training approaches, we tested whether models predicted fear in held out data in a situation-general manner across all three situations, or whether it predicted fear in a situation-dependent manner (i.e., in only one or two situations.

### Across-Situation Model Training: Shared Neural Codes

To determine whether there are brain regions wherein the same model parameters predict fear across situations, we trained the searchlight MVPA using data across all three situations and tested how well the model predicted fear for every situation using held out data. If a brain region contains situation-general neural codes in predicting fear, then training the model using data across situations should predict fear across situations. Alternatively, if a brain region contains situation dependent neural codes, then even upon training with data across situations, the model may only predict fear in one or two situations. As in prior studies, fear-predictive functional activity was widely distributed throughout cortical and subcortical areas (Figure 2). A breakdown of the situation general map showed that 1.9% of voxel-wise neighborhoods met criteria of having model parameters that predicted fear across situations. Of the remaining voxel-wise neighborhoods, 48.2% predicted fear in one situation, whereas 49.9% predicted fear in two of three situations. The reported findings use FWE-corrected significance tests, however, the proportions and locations of voxels classified as situation general and situation dependent did not substantially change when using a more lenient threshold (see Extended Data Figure 2-1). The small proportion of voxel-wise neighborhoods that predicted fear across situations were located in the right posterior insula and the right superior temporal cortex (Figure 2, red). Yet, the overwhelming majority of brain regions that predicted fear did so in only one or two situations.

**Figure 2.**
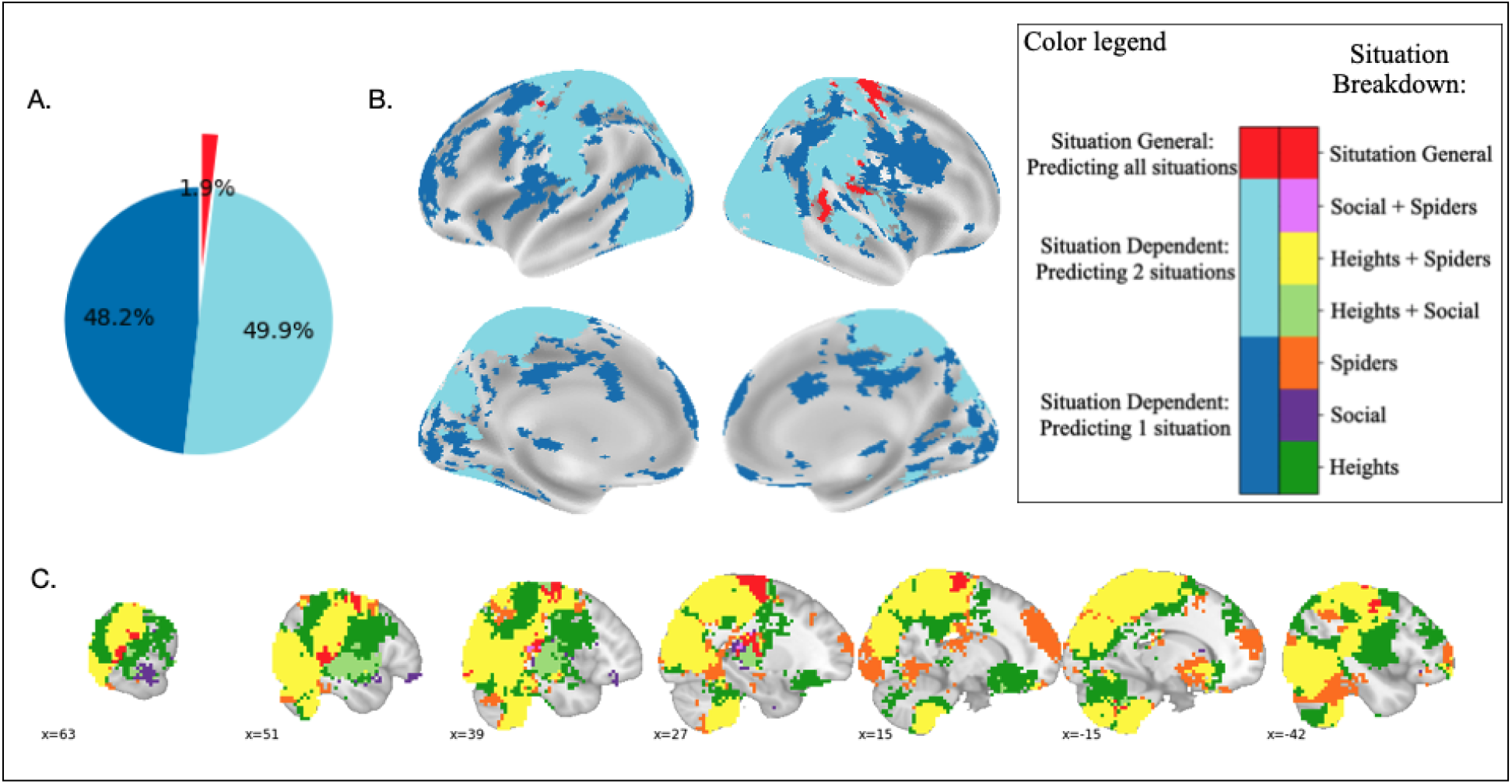
Across-situation model training: shared neural codes. Models were trained using data across all three situations. (A) The pie chart illustrates the percentage of voxels that predicted fear in held out data in one (dark blue), two (light blue), or all three (red) situations. Only 1.9% of voxelwise neighborhoods predicted fear across all three situations. (B) The maps illustrate which brain regions carried situation general and situation dependent information in predicting fear. Situation-dependent areas (both the dark blue and the light blue) are distributed across the whole brain. The situation-general areas (red) included the superior temporal cortex, posterior insula areas, and the somatosensory cortex. (C) A detailed breakdown of the neural pattern by each situation and their combinations. Color codes signify the specific situation or combination of situations predicted by the voxelwise neighborhood. Note, even with across-situation training, some areas only predicted fear for spider, social, or heights related stimuli.

### Situation-by-Situation Model Training: Unshared Neural Codes

To determine whether there are brain regions that predict fear reports across all situations, but that may use different neural codes depending on the situation, we trained and tested the searchlight with LASSO-PCR models situation by situation (i.e., trained and tested using data from the heights condition only, and the same for spider and social conditions). This approach resulted in a Pearson correlation map per situation. We performed a conjunction analysis (Nichols et al., 2005) to identify which voxel-wise neighborhoods predict fear across situations. This is a more lenient approach for identifying brain regions that predict fear across situations since the model parameters are allowed to vary by situation. Even so, a breakdown of the conjunction map showed that only 4% of fear-predictive voxel-wise neighborhoods carried information in all three situations (Figure 3). Of the remaining voxel-wise neighborhoods, 66.4% predicted fear in only one situation, and 29.5% predicted fear in two of the three situations. Notably, the nominal proportion of voxels that predicted fear in at least two situations doubled when using across-situation training (62%) relative to situation-by-situation training (30.5%), suggesting that the training type had an influence. However, if anything, across-situation training led to a nominal decrease in the proportion of voxels that contained situation general codes (i.e. from 4% to 1.9%). We present our findings using FWE corrected significance tests, however, we performed analyses across a range of lenient and stringent statistical thresholds to ensure that conclusions were robust across thresholding. The locations and proportions of voxel-wise neighborhoods classified as situation general or situation dependent did not meaningfully change when using a more lenient threshold (see Extended Data Figure 3-1).

**Figure 3.**
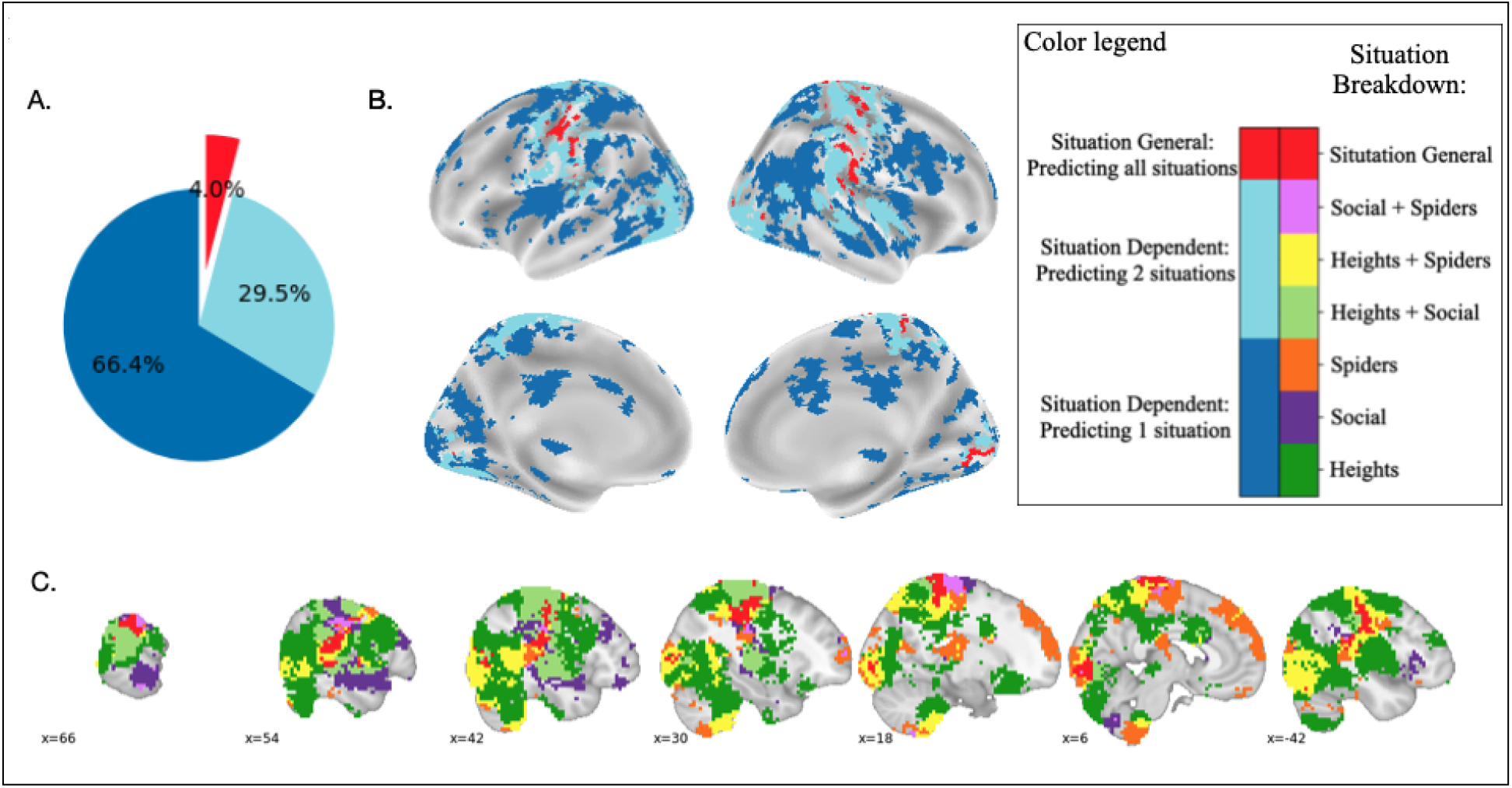
Situation-by-situation model training: unshared neural codes. Models were trained and tested on data from only a single situation, resulting in three maps. A conjunction analysis was used to identify which brain regions carried information that predicted fear across situations. This is a more lenient approach than across-situation training since model parameters are allowed to vary, situation-by-situation, in predicting fear. (A) The pie chart illustrates the percentage of voxels that predicted fear in held out data in one (dark blue), two (light blue), or all three (red) situations. 4% of voxelwise neighborhoods predicted fear across all three situations. (B) The maps illustrate which brain regions carry situation general and situation dependent information in predicting fear. The situation-general areas (red) included the superior temporal cortex and posterior insula areas, as before, but note these areas may involve model parameters that still vary by situation. (C) A detailed breakdown of the neural pattern by each situation and their combinations. Color codes signify the specific situation or combination of situations predicted by the voxelwise neighborhood.

## Discussion

In this study, we examined the extent to which the regionwise relationship between fear experience and neural activity reflects the functioning of a singular functional pattern or involves diverse, functionally heterogeneous brain states. We characterized each brain region based on whether it contained functional activity that predicted fear ratings across situations using either the same neural code (i.e. situation general, shared parameters) or flexible neural codes (i.e., situation general, unshared parameters), or alternatively, whether it only predicted fear in some but not all situations (i.e., situation dependent). For the overwhelming majority of brain regions, models of functional activity for predicting fear were situation dependent (98%). A small portion of voxel-wise neighborhoods (2%) predicted fear across all three situations using the same model parameters. Even upon allowing the model parameters to flexibly vary by situation, few areas (4%) carried information that predicted fear across all three situations. Overall, these results suggest that regional representations of fear are dominated by functionally heterogeneous, situation-dependent signals. This conclusion has im-portant implications for understanding both the neural representations and brain signatures of fear. The term “brain signature” may seem to imply uniformity of representation, and indeed, it has been interpreted as such in prior work. However, algorithms commonly used to estimate brain signatures can classify instances into emotion categories even when the underlying representational distribution of instances is degenerate (for details, see Clark-Polner et al., 2017; Kragel et al., 2018; Lindquist et al., 2022). Correspondingly, we suggest that “brain signatures” be viewed as an analytical approach wherein brain data is used to optimally predict behavior, but for which additional considerations are required to test theories regarding the neural representations of emotion (for example, see, Čeko et al., 2022). Questions regarding how the brain represents emotions lie at the crux of emotion theory (Barrett & Satpute, 2019; Ekman, 1992; Lindquist et al., 2013; Lindquist & Barrett, 2008; Mobbs et al., 2019; Panksepp, 2011). Our study was inspired by a constructionist approach, which posits substantial within-category heterogeneity in neural representations of emotion (Barrett, 2006, 2017b, 2017a; Doyle et al., 2022; Lindquist et al., 2012; Wilson-Mendenhall et al., 2015). According to this view, fear refers to a population category constituted from instances with diverse and heterogeneous features (Barrett, 2017b, 2017a; Siegel et al., 2018). Fear occurs when incoming sensory input is made meaningful with respect to similar previous instances in a predictive processing neural architecture (Barrett, 2017b). We reasoned that instances from similar situations are more likely to share features in common with each other, and thus may provide a useful heuristic to guide how instances serve as priors for conceptualizing future sensory inputs as yet another instance of fear (Satpute & Lindquist, 2019). By this account, fearful situations are not necessarily organized into “types” (e.g., a predator type, a heights type) with type-specific brain states. Rather, some instances of fear involving spiders, for example, may be similar to those involving heights, depending on the constituent features (Barrett, 2013; McVeigh et al., 2023). Consistent with this notion, a substantial portion of brain regions contained fear-predictive codes that generalized across two situations even though few brain regions predicted fear across all three situations. These findings coincide with recent theoretical and empirical approaches wherein context is integral to representation rather than modulating a core response profile (Satpute & Lindquist, 2019; Skerry & Saxe, 2015; Tamir et al., 2016; Wilson-Mendenhall et al., 2011). Appraisal theories propose that emotions are the outcomes of evaluations of the significance of an event in relation to one’s well-being and goals (Lazarus, 1991; Moors et al., 2013). If fear involves a particular appraisal configuration (Roseman & Smith, 2001) and specific appraisal dimensions involve the functioning of specific neural circuits or networks (e.g., amygdala for relevance appraisals, hippocampus/amygdala for novelty appraisals) (Brosch & Sander, 2013; Smith & Lane, 2015), then one might expect activity in those circuits to generalize in predicting fear across situations. Alternatively, more flexible appraisal models have proposed that instances of fear are associated with heterogeneous sets of appraisal patterns (Meuleman & Scherer, 2013) wherein appraisals are not necessarily causal antecedents of a “core fear” state, but rather are descriptive features of emotion (Ellsworth & Scherer, 2003; Ortony & Clore, 2015). Our findings showing substantial functional heterogeneity in fear may be more consistent with the latter than the former set of models. Yet, future work is required to determine whether the regional activation patterns are better explained by appraisal dimensions, or rather assembly representations of instances in line with the constructionist view. Functionalist models posit that fear refers to a goal (e.g., prevent harm from a predator) that may be achieved by different defensive behaviors (e.g., running, freezing, fighting; (Anderson & Adolphs, 2014; Fanselow, 1994; Fendt & Fanselow, 1999; Mobbs et al., 2019). These behaviors are thought to involve a circuit that traverses the amygdala, hypothalamus, and periaqueductal gray, among other primarily subcortical structures. Different configurations of this circuit may drive different defensive behaviors, depending on the situation (e.g., the imminence of the predator). It remains contested as to whether this circuit underlies both defensive behaviors and fearful experiences in a one-system model (Panksepp, 2011; Panksepp et al., 2011), or whether survival behaviors and fearful experiences involve distinct neural systems in a two-system model (LeDoux & Brown, 2017; LeDoux & Pine, 2016). Notwithstanding issues of spatial resolution with standard 3T fMRI (Satpute et al., 2013) and constraints of the searchlight approach (Kragel et al. 2018, Zhou et al. 2021), our findings suggest that single-system accounts may not fully account for “fear” (Taschereau-Dumouchel et al., 2020, Kragel & LaBar 2015). For instance, functional activity in the amygdala predicted fear in some, but not all, situations – even when the neural codes were allowed to vary to accommodate the idea that a single circuit may engage in different functional configurations to support fear. Establishing boundary conditions for generalization would be a critical avenue for future work. For instance, macrolevel architecture associated with defensive behavior may generalize in predicting fear in situations that share features of a predator-prey interaction, such as predatory imminence (Fanselow, 1994), or alternatively, when there are similar allostatic demands, regardless of whether the situation resembles predator-prey (Barrett & Finlay, 2018; Rosen & Schulkin, 2004). Notably, functional activity in some brain regions, including the temporal-parietal area and posterior insula, may support situation-general representations. Areas overlapping with (Skerry & Saxe, 2015) or contralateral to (Peelen et al., 2010) the posterior superior temporal sulcus have been implicated in emotion categorization but in the context of emotion perception. The posterior insula receives sensory inputs from the body and may play a more general role in arousal or interoception that is shared across mental phenomena (Carvalho & Damasio, 2021; Craig, 2002, 2009; Damasio, 1999; Kleckner et al., 2017; Satpute et al., 2019). However, these areas have also been inconsistently implicated in prior MVPA studies on emotion experience (Extended Data Table 1-1). Future work using other stimulus modalities (e.g., fear of pain) and assaying other emotions is required to test whether these areas carry generalizable, and specific, neural codes that predict fear. Our findings underscore the importance of testing for external validity and generalizability of a given brain-behavior relationship (Lee et al., 2021; Shackman & Wager, 2019). Many studies in affective neuroscience preclude tests for external validity by examining fear in a single context or averaging findings across trials. Yet, our findings suggest that generalizability may be strongly constrained by the situation. To effect, and perhaps owing to the lack of robust predictive models of valence (for a review, see Lee et al., 2021), recent theoretical models in affective neuroscience incorporated modality as an organizing factor (Chang et al., 2015; Chikazoe et al., 2014; H.-C. Kim et al., 2019; J. Kim et al., 2017; Lee et al., 2021; Miskovic & Anderson, 2018; Satpute et al., 2015). For instance, recent work has advanced a “visually induced fear signature” (Zhou et al., 2021). Yet, we only used visual stimuli and yet we still found robust evidence of context-dependence. Thus, our findings suggest that representations of emotion categories are not necessarily organized into modality dependent “types”, but rather that the sensory modality is just one aspect of a broader interpretation of context, wherein context might be better characterized in terms of predictions and prediction errors that are derived from prior experience (Barrett, 2022; Lee et al., 2021). Insofar as fear lies at the crux of emotion theory, it stands to reason that other emotion categories, too, are likely to exhibit degeneracy, or many-to-one functional mappings between brain states and psychological constructs (Barrett & Satpute, 2019; Doyle et al., 2022; Friston & Price, 2003; Khan & Wang et al., 2022). Notably, our findings converge with recent work showing strong evidence of situation dependence in the peripheral autonomic correlates of fear, too (McVeigh et al., 2023). Modeling this variation may be key to developing a fundamental understanding of complex mind-brain-behavior relationships alongside personalized treatments in clinical populations.

## Data sharing plan

Anonymized data will be deposited in OpenNeuro (https://openneuro.org/) after publication. Analysis scripts are available in Github at https://github.com/yiyuwang/AffVids_mvpa

## Author Contribution

A.B.S designed the study and collected the data; Y.W., P.A.K., and A.B.S. contributed analytic ideas; Y.W. analyzed the data; Y.W., P.A.K., and A.B.S. wrote the manuscript.

