## Extended Data for "Neural predictors of subjective fear depend on the situation"

**Extended Data: Neural predictors of subjective fear depend on the situation**

**Stimulus Norms (supporting information for Figure 1).** Participants (N = 100) recruited from Amazon Mechanical Turk rated video stimuli on fear, valence, and arousal using 7-point likert scales (1 = "low fear", 7 = "high fear"). Normative ratings were resampled to the range of 0 and 1, and were used to select stimuli that vary from low to high levels of normative fear within each content condition (social, spiders, heights). However, our research question and primary analyses focused on fear ratings that were provided by each participant who completed the fMRI. Descriptive states and video descriptions are available at: https://github.com/yiyuwang/AffVids_mvpa/tree/main/video_info

**Interpolation (supporting information for Figure 1)**. Participants failed to provide fear ratings within the allotted time on a small proportion of trials (mean missing trials per person = 1.23, max = 8). Missing fear ratings were interpolated using normative ratings by substituting the normative ratings but rescaled to fit the individual's range of the fear ratings of the specific situation. Of note, recalculating analyses upon excluding missing data did not significantly alter the findings. Fear ratings were z-scored within each situation for each participant, to remove idiosyncratic differences in rating scale interpretations by participant while maintaining fidelity to subjective variation in fear across stimuli. We used the same interpolation procedure for arousal and valence ratings, but our analysis did not examine arousal and valence and they were not used in this manuscript.

Figure 1-1. Subject fear ratings by video order

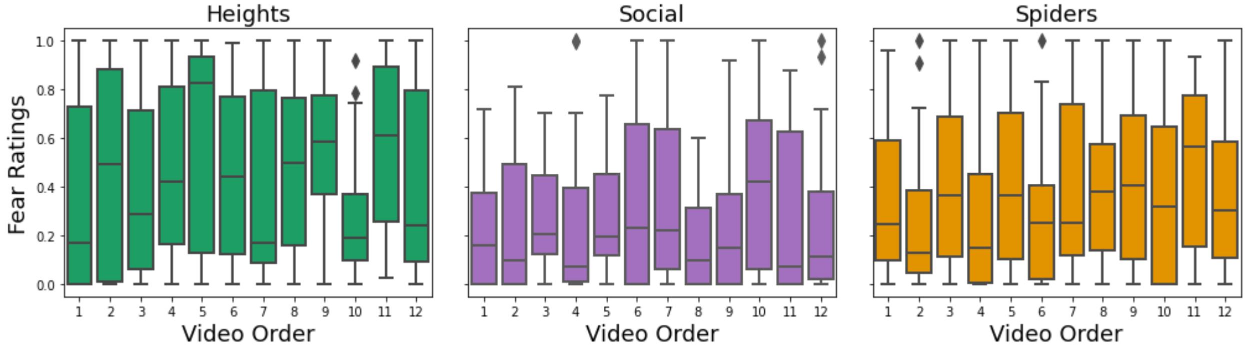


Figure 1-1. Self-reported fear ratings did not show sensitization or habituation effects over time for each situation. Participants watched a total of thirty six videos across three functional runs in the fMRI scanner, such that each run consists of four videos from each of the three situations. The box and whisker plots show mean and distribution of fear ratings across videos, rank ordered across time.

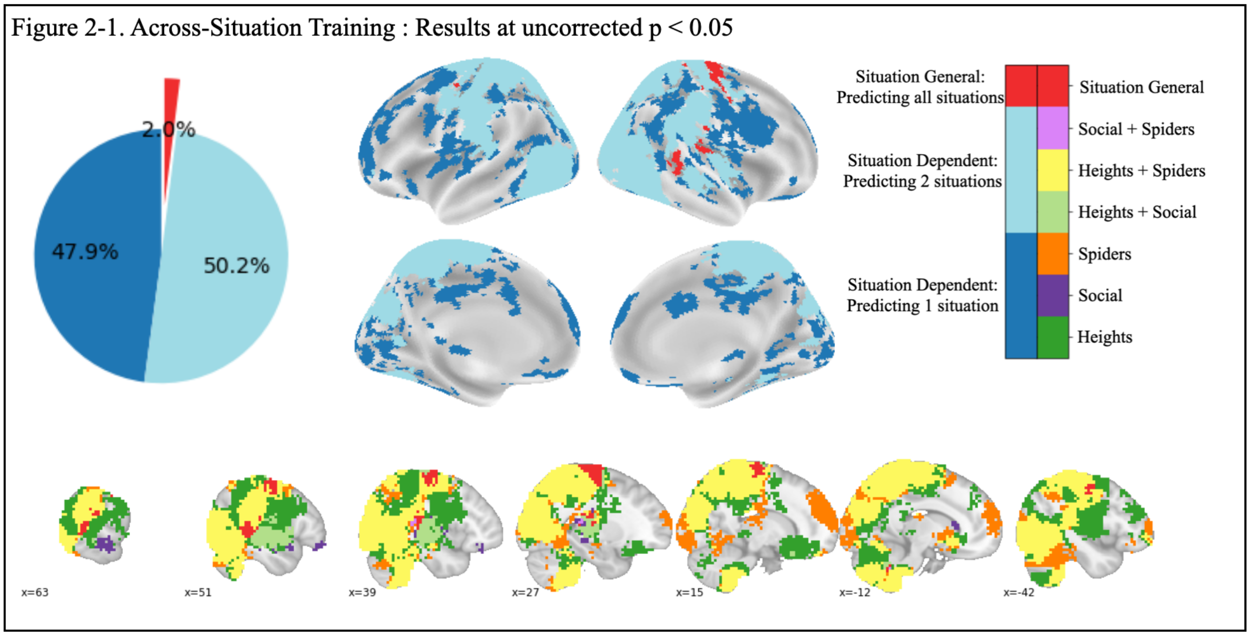


Figure 2-1. Results from the Across-Situation training using uncorrected $p < 0.05$: The proportion of voxels classified as situation general and situation dependent or their general locations did not significantly change when using a more lenient threshold.

Figure 3-1. Situation-by-Situation Training: Results at uncorrected $p < 0.05$

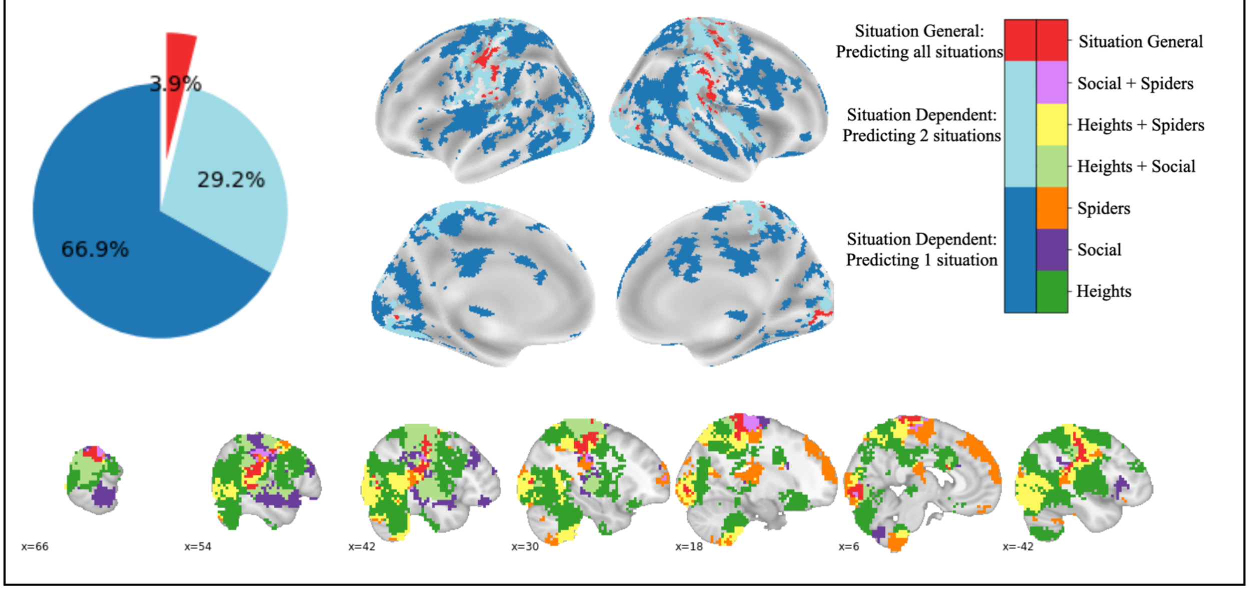


Figure 3-1. Results from the situation-by-situation training using uncorrected $p < 0.05$: The proportion of voxels classified as situation general and situation dependent or their general locations did not significantly change when using a more lenient threshold.

Figure 3-2. Situation General Model Sample Size Comparison

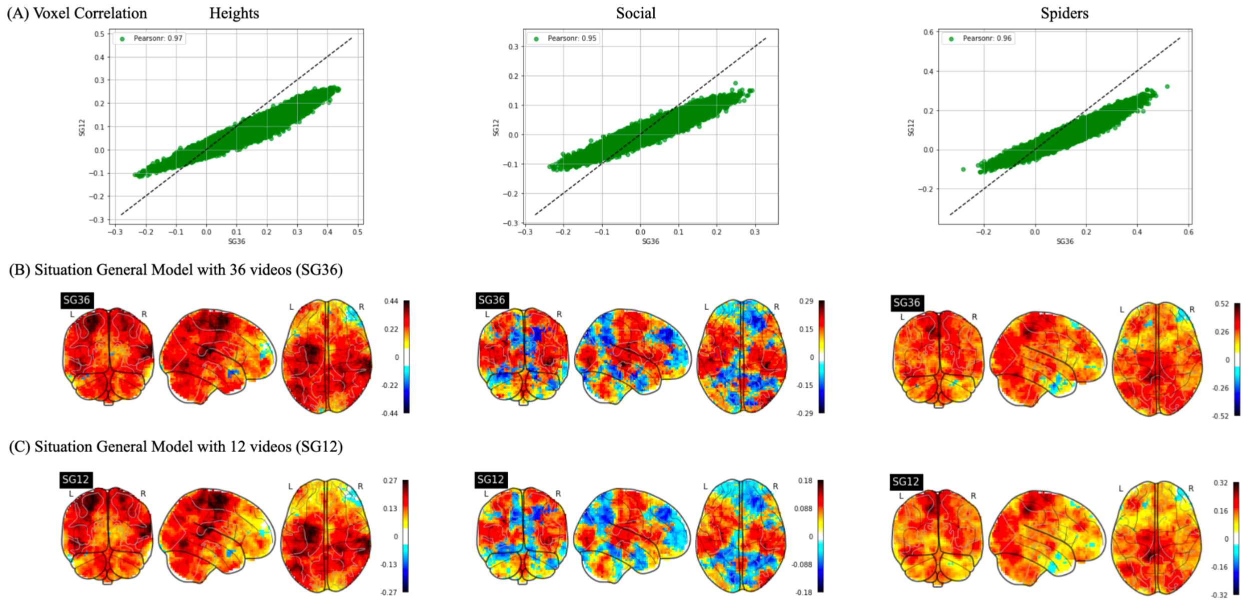


Figure 3-2. Correlation between different numbers of training samples in the across-situation model. The training schemes in the situation-by-situation training strategy included 12 samples, while the across-situation training strategy included 36 videos("SG36"). To make sure that the difference between the sample sizes of the two training strategies did not influence the results, we performed a random sub-sampling and compared the results with the full sample results. First, twelve videos were randomly sampled, four from each situation as the training sample. We repeated the random sub-sampling and training 200 times and calculated an average model ("SG12"). (A) The correlation values between the SG12 and SG36 are 0.97, 0.95, 0.96 for the situation of Heights, Social, and Spiders respectively, suggesting that the patterns are comparable between SG12 and SG36. (B-C) Whole brain patterns for SG36 and SG12. Noticeably, and as might be expected, the r values are lower in the SG12 than in the SG36, which could be due to fewer shared video stimuli for SG12 with the testing sample. Only four videos could be shared between the training and the testing sample in SG12, whereas both the SG36 and the situation-dependent training strategy have all twelve video stimuli shared between the training and the testing sample. Since our hypothesis is not about comparing the model performance between the situation-dependent and the situation-general model, it does not make sense to present a situation-general model with a reduced number of training samples to match the other model. As such, we decided to keep the situation-general model that included all the data. Future studies with more extensive sample size could investigate the relationship between training sample size and effect size.

Figure 3-3 Signal Check for OFC and Amygdala: functional activity

(A) Medial OFC, slices at [2, 38, -16]

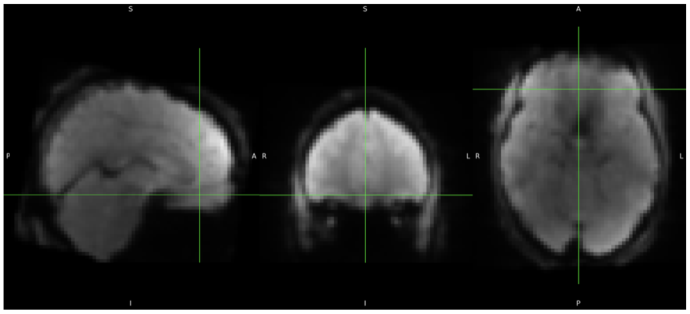

(B) Amygdala, slices at [23, -7, -13]

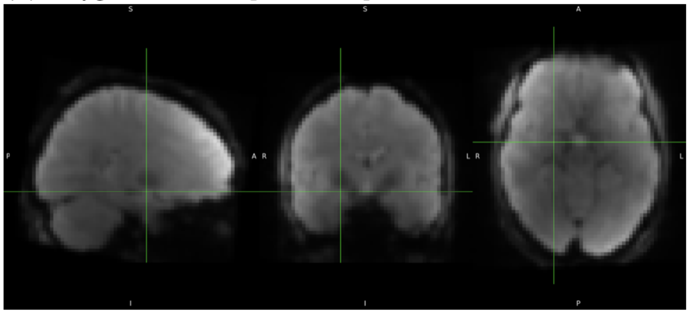

Figure 3-3. The figure presents a sample of time-averaged EPI scan for assessing coverage. The cursor is centered on the medial OFC (panel A), and the amygdala (panel B). The sequence captured functional activity in both regions.

Figure 3-4. Signal Check for OFC and the Amygdala: Time Series

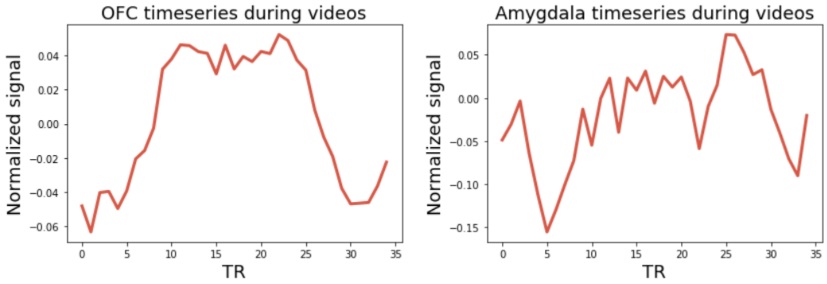


Figure 3-4. Time series during video watching period, averaged over participants and all videos. 0 TR indicates the video onset. Time series suggest that functional activity was captured.

| ROI name | Voxels | Center of Mass (mm) | | | Max Pearson's r value | | |
|---|---|---|---|---|---|---|---|
| | | X | Y | Z | Heights | Social | Spiders |
| Primary motor cortex | 419 | 30.1 | -15.5 | 65 | 0.387 | 0.268 | 0.338 |
| Superior temporal cortex* | 247 | 42.7 | -30.8 | 11 | 0.41 | 0.274 | 0.354 |
| Premotor cortex | 42 | -45.1 | -9.93 | 51 | 0.405 | 0.209 | 0.279 |
| Somatosensory cortex | 23 | 63.1 | -25.2 | 23 | 0.405 | 0.206 | 0.255 |
| Cerebellum | 9 | -10.7 | -55.3 | -38 | 0.305 | 0.225 | 0.21 |
| Visual cortex | 6 | 40 | -88 | -11.5 | 0.372 | 0.187 | 0.313 |
| Hippocampus/Insula | 6 | 35 | -14.5 | -9.5 | 0.29 | 0.244 | 0.232 |
| Superior temporal cortex | 5 | 46.2 | -11.4 | -11.4 | 0.293 | 0.211 | 0.215 |

*The center of mass located in the superior temporal cortex, but the activation cluster included both the superior temporal area and the Insular area.

Table 2-1. Situation general ROIs from the Across-situation training
ROIs and its center of mass for the situation general areas in the Across-situation training (red areas in Figure 2)

Table 2-2. MVPA studies of fear signatures

| Study | Analytical Algorithm (emotion classification OR fear degree) | Searchlight vs ROI vs Whole brain | Stimulus Types | Generalization | Right posterior lateral temporal cortex | Posterior insula |
|---|---|---|---|---|---|---|
| 2010 Peelen et al. | Emotion Category Classification | ROI | Pictures of Faces/Bodies | Yes (faces/bodies) | No (left STS) | No |
| 2015 Kragel and LaBar | Emotion Category Classification | Whole Brain | Music/Film | Yes (music/film) | No | No |
| 2015 Wager et al. | Emotion Category Classification | Whole Brain | Meta Analysis | No (diverse studies, but not systematic) | No | No |
| 2015 Skerry and Saxe | Emotion Category Classification | Searchlight + Whole Brain | Verbal descriptions of scenarios | No (diverse stimuli, but not systematic) | Adjacent, not overlapping | No |
| 2016 Saarimaki et al. | Emotion Category Classification | Whole Brain | Music/Film | Yes (music/film) | No | Yes |
| 2018 Saarimaki et al. | Emotion Category Classification | Whole Brain | Auditory narratives of scenarios | No (diverse stimuli, but not systematic) | No (deactivation) | Yes |
| 2020 Taschereau-Dumouchel et al. | Subjective Fear | ROI | Pictures of animal categories | No (diverse stimuli, but not systematic) | Yes | No (left insula) |
| 2021 Zhou et al. | Subjective Fear | ROI+ Whole Brain | IAPS | No (diverse stimuli, but not systematic) | Adjacent, not overlapping | Negative correlation |

| ROI name | Voxels | Center of Mass (mm) | | | Max Pearson's r values | | |
|---|---|---|---|---|---|---|---|
| | | X | Y | Z | Heights | Social | Spiders |
| Primary motor cortex | 759 | 37.5 | -26.8 | 45.6 | 0.43 | 0.279 | 0.356 |
| Somatosensory cortex | 408 | -51.5 | -21.9 | 34.5 | 0.371 | 0.28 | 0.352 |
| Occipital cortex | 128 | 7.57 | -87.6 | -3.7 | 0.288 | 0.262 | 0.385 |
| Occipital cortex | 54 | 37.6 | -80.9 | 5.7 | 0.389 | 0.224 | 0.308 |
| Temporal parietal cortex | 30 | -56.9 | -64 | 16.5 | 0.289 | 0.245 | 0.333 |
| Superior temporal cortex | 5 | -54.9 | -40.2 | 6 | 0.233 | 0.229 | 0.282 |
| Motor cortex | 5 | -54.6 | 1.8 | 11 | 0.216 | 0.218 | 0.206 |

Table 3-1. Situation general ROIs from Situation-by-situation training
ROIs and its center of mass for the situation general areas in the Situation-by-situation training (red areas in Figure 3)
